# Impact of Species-Specific Plant Growth-Promoting Rhizobacteria (PGPR) on Maize (*Zea mays*) Phenotypic and Biochemical Diversity

**DOI:** 10.1101/2024.04.28.591576

**Authors:** Swapnil Singh, Rajib Roychowdhury, Arkadeep Mukherjee, Harleen Kaur, Ravneet Kaur, Neetu Jagota, Surinder Sandhu, Vinod Kumar, Mahiti Gupta, Young-Ho Ahn, Vineet Meshram, Ashish Sharma

## Abstract

Maize (*Zea mays*) is a vital cereal crop used as a staple diet in many countries. In contemporary farming practises, inoculation with plant-growth-promoting rhizobacteria (PGPR) can be promoted as a sustainable alternative to chemical fertilizers and pesticides in cereal crops including maize. For experimental verification of the above-mentioned hypothesis, four PGPR inoculants *Pseudomonas fluorescens*, *P. putida, Azospirillum lipoferum* and *Bacillus subtilis* were applied to three maize genotypes (AXE*, PMH1 and PMH10) and their effects were studied by measurement of various morphological and biochemical paramters. Substantial increase in the amount of chlorophyll a (45%), chlorophyll b (100%), total chlorophyll (95%), nitrate reductase (22%), superoxide dismutase (52%), protein content (16%), methionine content (31.8%), grain protein content (69%) were recorded over the control (non-treated or mock) plants. Morphological parameters also exhibited an increase in leaf number (53%), fresh weight (41%) and dry weight (62%) in test plants. Comparable outcome was observed for whole cob weight showing an increase of 42%, cob weight (60%), hundred-grain weight (25.9%), maize grain yield m^-2^ (18%) and yield ha^-1^ (18%) over the control. This study highlights the efficacy of the aforementioned four PGPR species as the most effective for maize crops. *Bacillus subtilis* and *Azospirillum lipoferum* may be considered species-specific PGPRs due to their superior performance compared to other strains. The considerable growth-promoting advantages observed in maize plants treated with bacterial inoculation indicated that PGPRs have the potential to be employed as sustainable solutions in maize production.

## 1 Introduction

Among the cereals, maize has the largest genetic yield potential. All plant parts of maize have trade and industrial value and can be used to make a wide range of foods. The crop has wide adaptability and can be grown in a variety of climates ranging from sea level to 3000 m above sea level (Sood et al., 2020). Maize serves as the principal raw material for many different manufacturing commodities as well as primary meal and nutritious feed for livestock. The agricultural sector, vital for the survival and sustenance of a nation, has experienced a surge in productivity over the past decades, thanks to technological advancements. However, these innovations have led to environmental challenges (Basu et al., 2021). To manage these challenges along with the assurance of high agricultural productivity, shifting focus towards biological interventions can be an effective strategy (Liu et al., 2021a). To this effect use of plant-based extracts and vermiculite teas can be effective in replacing destructive agrochemicals and maintaining soil health (Arancon et al., 2019). Effective plant growth-promoting rhizobacteria (PGPR) are currently examined as organic fertilizers and biological pest controllers to minimize the consumption of artificial agricultural chemicals in the farming industry (Anli et al., 2020). Optimistic impacts of PGPR on both quantitative and qualitative plant attributes have been well documented in maize. Pérez-Pérez et al. (2021) reported positive effects of *Stenotrophomonas*, *Pseudomonas* and *Rhizobium* on maize growth. Singh et al. (2023) showed synergistic effects of various PGPR on the growth promotion of maize. Plant biomass, root parameters and nutrient content of maize plants can be improved significantly by the inoculation of PGPR (Song et al., 2023). An increase in the contents of nutrients, specifically iron and zinc has also been reported in response to the inoculation of PGPR from *Bacillus* and *Paenibacillus* species (Ahmed et al., 2020). Inoculation of maize plants with *Cupriavidus necator* and *Pseudomonas fluorescens* improved root/shoot ratio by almost 40% and aerial biomass by approximately 89% in the inoculated plants under water stress (Pereira et al., 2020). Separate inoculation of the PGPR strains *Sinorhizobium, Bacillus, Sphingomonas* and *Enterobacter* to the maize rhizosphere significantly improved maize growth and grain yield (Chen et al., 2021).

Plant growth-promoting rhizobacteria has been shown to improve nutrient quality while minimising crop growth risks. According to Kumar et al. (2018) Plant growth is dependent on chlorophyll and nitrate reductase activity. To evaluate nitrate assimilation in plants, nitrate reductase assays were employed and chlorophyll content was examined to monitor the state of the plants, PGPR inoculation improved both leaf nitrate reductase and chlorophyll content as compared to their respective controls. Nawaz and Bano (2020) reported, antioxidant enzymes for example superoxide dismutase (SOD) protect plants from any harm initiated by oxidative stress, plant cells contain SOD, which detoxifies reactive oxygen species (ROS) and protects plants from oxidative stress. Superoxide dismutase activity increased in *Phaseolus coccineus* commonly known as runner bean plants treated with *Pseudomonas putida*. The build-up of osmolytes such as proteins in plants aids in maintaining the water balance of cells, and sub-cellular structures as well as the resistance for membranes along with other proteins against stress caused by osmotic pressure (Naseem et al., 2018). Carbohydrate was recognised to operate as an osmoprotectant, that stabilises cellular membranes and sustains plant turgor. Sucrose and hexoses regulate gene expression and affect the expression of genes linked to stress. Soluble carbohydrates constitute key osmolytes in plants which help in osmotic adjustment during stressful circumstances. Bacterial inoculation with *Bacillus* spp. and *Pseudomonas* spp. greatly increases protein and carbohydrate content while also protecting against various abiotic stresses (Naseem et al., 2018). Methionine plays a crucial role in sulphur synthesis and is essential for the production of glutathione, a key antioxidant enzyme that inhibits free radical activity. Methionine deficiency not only hampers growth but also has an effect on sulphur metabolic pathways since the methionine structure contains sulphur (Pandey et al., 2018). Phytic acid, which typically occurs in Fabaceae and Gramineae families, has been shown to reduce iron and zinc bioavailability in plant-based meals and catalase is an enzyme found in all living beings that are aerobic that catalyses H_2_O_2_ into O_2_ and H_2_O, thereby shielding cells from oxidative deterioration triggered by ROS (Uzma et al., 2022). The majority of the damage occurs at the cellular level as a consequence of oxidative damage triggered by drought stress; this injury is produced by a disproportion between the construction of ROS and their purification, inoculation with PGPR lowers anti-nutrient activity in addition to abiotic stress (Khan et al., 2019).

Hence, the current investigation was conducted to determine the impact of PGPR inoculation on the biochemical activities of the maize plant based on the hypothesis that PGPR can aid in plant growth. The primary goal of this study is to determine the species-specific PGPR for maize genotypes.

## 2. Materials and methods

### 2.1 PGPR isolates procurement and maintenance

Plant growth-promoting rhizobacterial inoculants of *Azospirillum lipoferum* (MTCC2694)*, Bacillus subtilis* (MTCC121)*, Pseudomonas fluorescens* (MTCC103) and *P. putida* (MTCC102) were procured from Microbial type culture collection (MTCC), Institute of Microbial Technology, Chandigarh, India, and used in this present study. The inoculants were cultured in Luria Broth (LB) medium with the cultures kept in a shaking incubator at 150 rpm and 29°C for 24 hours. Their OD was adjusted to 1 spectrophotometrically (λ=660 nm) which corresponds to a count of approximately 10^6^ cfu for inoculation. For storage purposes, the cultures were inoculated to a solid LB medium and kept at 4°C.

### 2.2 PGPR treatments to the plants

Commercially available maize genotype AXE*, PMH1 and PMH10 were used in this study and the seeds were procured from the Department of Plant Breeding and Genetics, Punjab Agricultural University (Ludhiana, Punjab, India). The seeds were surface sterilized and sown in the experimental fields of the Faculty of Agricultural Science, DAV University, Jalandhar, where plants were 25 cm apart in a single row with a spacing of 75 cm amongst different rows with 4 rows per plot, giving it the dimensions of 3 m × 2 m and a total of 32 plants per plot. In total, 45 such plots were sown (supplementary figure 1). Inoculation of the seedlings was done at 3 leaf stage, for inoculation each seedling received 20 ml of bacterial culture in the form of a spray to the soil (Moreira et al., 2014). Irrigation and application of basal fertilizer dose in the form of Diammonium Phosphate (DAP) was done as per recommendations of Indian Council for Agricultural Research (ICAR), New Delhi, India for maize growth. Along with the treated plants untreated controls were maintained for each genotype.

### 2.3 Phenotypic evaluation

Flag leaf and penultimate leaf were used for recording observations pertaining to biochemical parameters while for leaf number, plant fresh and dry weight, and yield data above ground plant biomass was considered for the plants under study. The observations were recorded at the vegetative stage which falls roughly 4 weeks after sowing (after 2 weeks of inoculation) and at panicle emergence (designated as flowering stage) which falls roughly 8 weeks after sowing (after 6 weeks of inoculation). Three replicates were selected from each plot at random for the measurement of various parameters. The experiment culminated in the harvesting of the crop approximately 90 days after the initial sowing along with the documentation of data at and after harvesting.

### 2.4 Morphological parameters

Fresh weight of above ground plant biomass, weight of covered and naked cobs, was determined immediately after the plants were harvested (90 days after initial sowing) from the plot. The plants were dried at room temperature for 6 days to get a consistent weight for determining dry weight (g). The leaf number was determined at regular intervals. Following harvest, grain weight and yield were also recorded.

### 2.5 Biochemical parameters

#### 2.5.1 Nitrate reductase (NR) activity

The method suggested by (Hageman and Hucklesby, 1971) was used to determine NR activity in the test and control plants. For measurement, 0.5 g of finely chopped fresh leaves were incubated with an equal ratio mixture of ice-cold phosphate buffer (0.02M, pH 7) and KNO_3_ at 33 °C for 30 minutes away from light. After that, 0.2 ml of the incubated solution was mixed with 0.5 ml each of 1% sulphanilamide and 0.5% NEDD [N-(1- Naphthyl)ethylenediamine] and the colour was allowed to develop and its absorbance was recorded at 540 nm for calculation of NR activity.

#### 2.5.2 Chlorophyll content

Chlorophyll quantification was done by the method given by (Hiscox and Israelstam, 1979). Finely chopped fresh leaf sample (50 mg) was mixed with 10 ml DMSO and incubated at 65°C for 30 min with periodic shaking. The absorbance of clear supernatant was recorded at 663 and 645 nm and calculations were made according to the formula described in Singh et al. (2023).

#### 2.5.3 Superoxide dismutase (SOD) activity

The SOD activity was assayed following the procedure outlined by Beauchamp and Fridovich (1971), and calculations of enzyme activity were done using the method of Giannopolitis and Ries (1977).

#### 2.5.4 Protein content

Protein content was assayed following Bradford (1976). For determination of protein content 0.2 g fresh leaf sample was crushed in Tris buffer followed by centrifugation at 8000 g for 15 min at 4 °C. After centrifugation 0.2 ml supernatant, 3 ml Bradford dye and 0.8 ml distilled water were mixed and absorbance was recorded at 595 nm.

#### 2.5.5 Carbohydrate content

The protocol given by Sadasivam (1996) was employed for the determination of carbohydrate content in grains. Crude seed extract was mixed with 6 ml Anthrone reagent in boiling tubes and placed in a boiling water bath for 10 min flowed by immediate cooling in an ice bath. The tubes were later incubated at 25 °C for 20 min for colour development. Absorbance was determined at 625 nm and the carbohydrate content was determined against the standard curve of sucrose.

#### 2.5.6 Catalase activity

The initial rate of disappearance of H_2_O_2_ has been employed to evaluate catalase (CAT) activity according to the method given by Kato and Shimizu (1987). The drop in H_2_O_2_ content at 240 nm in the reaction mixture containing 50 µl enzyme extract was recorded and CAT activity was determined using H_2_O_2_ extinction coefficient of (40 mM^-1^ cm^-1^ at 240 nm).

#### 2.5.7 Phytic acid

A modified version of Harland and Oberleas (1977) and Latta and Eskin (1980) protocol was used to extract phytate fromseeds of maize plants. 2.4 % HCl (0.65 N) was used for the initial extraction, because it was more effective at removing all of the phytate. The phytate concentration was assessed with the Wade reagent in addition with the digestion process at 500 nm by spectrophotometer.

#### 2.5.8 Methionine content

The protocol described by Horn et al. (1946) was used for the determination of the amount of methionine in grain. A yellow colour adduct is formed when the methionine released from tissues is mixed with nitroprusside solution at high pH which gradually turns into a red colour upon acidification. Glycine prevents the unwanted reaction of nitroprusside with other amino acids if present.

#### 2.6 Statistical analysis

The source of variation in the data obtained was analysed using SPSS16 statistical package. The parameters were subjected to ANOVA and Duncan’s two-sided test for comparison of treatments at the significance level of 0.05 (Cyprien and Kumar, 2012).

## 3. Results

### 3.1 Morphological parameters

A significant rise was recorded in leaf number, and fresh and dry weight of the maize plants during 2021 and 2022 in comparison to the control plants (Figure 1). During 2021, the number of leaves increased by 46 % for AXE*, 56% for PMH1, and 36% for PMH10 plants inoculated with *A. lipoferum, P*. *fluorescens* and *B. subtilis* respectively in comparison to the control plants. The fresh weight of the plant increased by 22% for AXE*, 24% for PMH1, and 21% for PMH 10. Inoculation with *A. lipoferum* resulted in maximum fresh weight for all the genotypes under study. The increase in dry weight of plant was observed by 53% for AXE*, 49% for PMH1, and 65% for PMH10, inoculated with *B. subtilis, P. putida* and *A. lipoferum,* respectively. Furthermore, during 2022, the rise in the number of leaves was 53% for AXE* and 31% for PMH1 inoculated with *B. subtilis*, and 34% for PMH10, inoculated with *A. lipoferum* and *B. subtilis* in comparison to the control plants. The increase in fresh weight of the plant for AXE* was 29%, for PMH1 was 41% and for PMH 10 was 34%. Inoculation with *A. lipoferum* for PMH1 and AXE* *B. subtilis* for PMH10 had the highest fresh weight. Plant dry weight increased by 62% for AXE*, 55% for PMH1, and 62% for PMH10, resulting in AXE*, PMH 1 and PMH10 plants inoculated with *A. lipoferum* having the greatest increase in plant dry weight.

**Figure 1.**
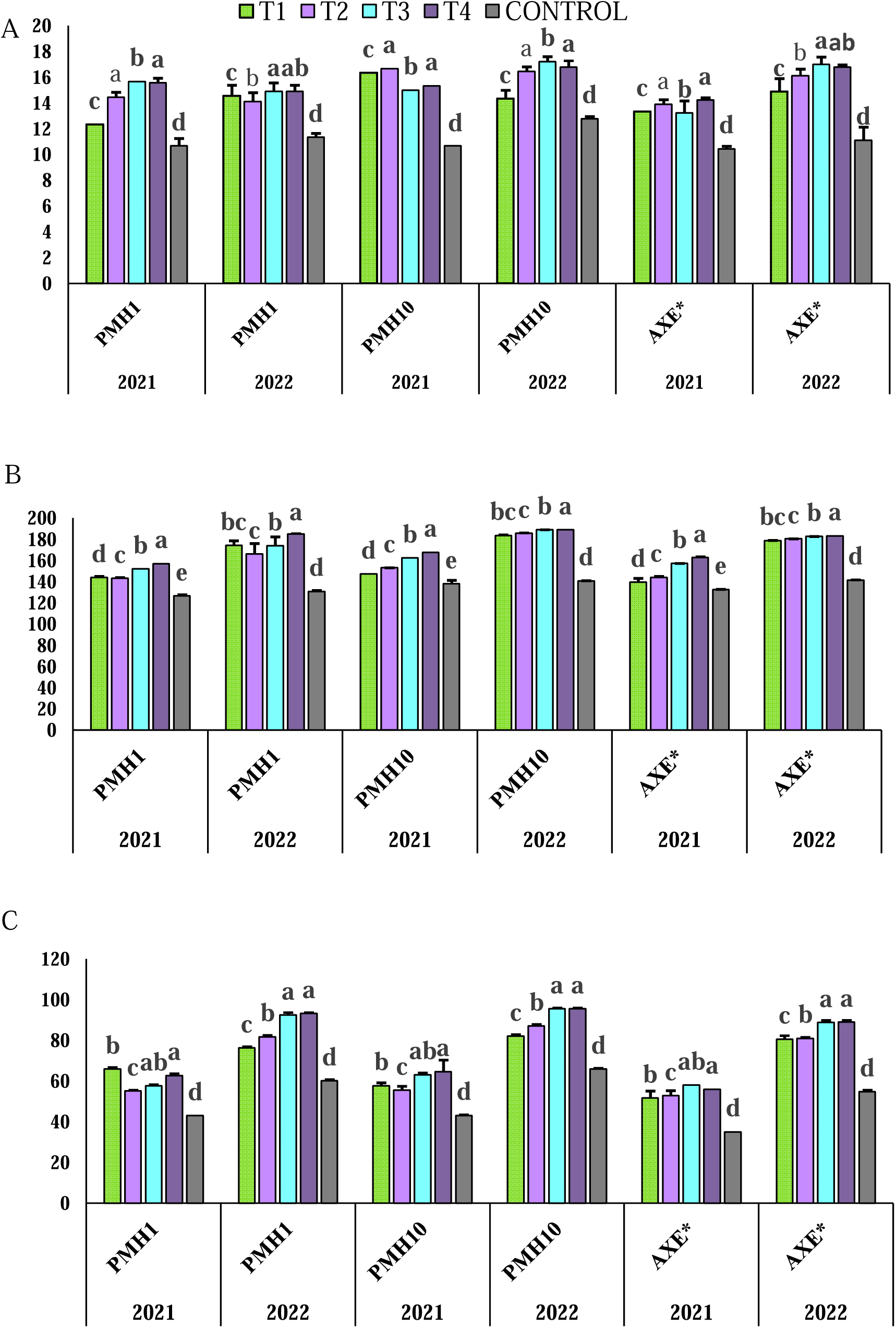
Effect of application of different plant growth promoting rhizobacteria on maize in three genotypes as morphological data (A) fresh weight of plant (B) dry weight of plant (C) number of leaves in years 2021 and 2022. Here T1 represents *P. putida (*MTCC 102), T2 represents *P. fluorescens* (MTCC 103), T3 represents *B. subtilis* (MTCC 121), T4 represents *A. lipoferum* (MTCC 2694). Means in the columns that have a common letter after them are not significantly different.

### 3.2 Biochemical parameters

Biochemical parameters viz. NR activity, SOD activity, protein content, chlorophyll a, chlorophyll b, total chlorophyll and CAT were also recorded in the present investigation to find out the general health of plants and the data obtained showed the following results at the vegetative stage and flowering stage.

#### 3.2.1 Nitrate reductase

During 2021 (Table 1) inoculation with different PGPR resulted in enhanced NR activity at the vegetative stage. In the genotype AXE* inoculation with *P. fluorescens,* resulted in the highest NR activity (1.98 Ug^-1^FWh^-1^) whereas values recorded for control was 0.96 Ug^-1^FWh^-1^, for PMH1 maximum NR activity was observed in the treatment with *P. putida* (4.83 Ug^-1^FWh^-1^), while the value for control was 0.95 Ug^-1^FWh^-1^ and for PMH10 highest NR activity was observed in the plants treated with *B. subtilis* (2.35 Ug^-1^FWh^-1^) against 0.95 Ug^-1^FWh^-1^ observed for control plants. However, during 2022 (Table 2), the highest rise for NR activity was observed in plants inoculated with *P. putida* for genotypes AXE*, PMH 1, and PMH10, with the observed values of 3.30 Ug^-1^FWh^-1^ against 0.94 Ug^-1^FWh^-1^ for control in AXE*, 3.71 Ug^-1^FWh^-1^ against 0.77 Ug^-1^FWh^-1^ for control in PMH1, and 3.23 Ug^-1^FWh^-1^ against 0.36 Ug^-1^FWh^-1^ for control of the genotype PMH10.

**Table 1.** Nitrate reductase (NR) activity (U g^-1^ FW h^-1^), Superoxide dismutase (SOD) activity (U g^-1^ FW h^-1^), Protein content (PR) (mg g^-1^FW), Catalase activity (CAT) (mg g^-1^ FW), Chlorophyll a content (Chl. a) (mg g^-1^ FW), Chlorophyll b content (Chl. b) (mg g^-1^ FW), Total Chlorophyll content (Total Chl.) (mg g^-1^ FW) in the leaves of maize genotype treated with different bacterial strains and untreated control in vegetative stage in 2021. The values after ± indicated deviation from the mean value. Means in the columns that have a common letter after them are not significantly different.

| Genotype | Treatment | NR<br>($\text{U g}^{-1} \text{FW h}^{-1}$) | SOD<br>($\text{U g}^{-1} \text{FW h}^{-1}$) | PR<br>($\text{mg g}^{-1} \text{FW}$) | CAT<br>($\text{mg g}^{-1} \text{FW}$) | Chl. a<br>($\text{mg g}^{-1} \text{FW}$) | Chl. b<br>($\text{mg g}^{-1} \text{FW}$) | Total Chl.<br>($\text{mg g}^{-1} \text{FW}$) |
| --- | --- | --- | --- | --- | --- | --- | --- | --- |
| PMH 1 | *P. putida* | 1.69±0.04<sup>b</sup> | 4.29±0.16<sup>c</sup> | 58.31±1.53<sup>b</sup> | 140.44 ±4.44<sup>b</sup> | 0.75±0.02<sup>a</sup> | 0.79±0.03<sup>a</sup> | 0.91±0.01<sup>a</sup> |
| PMH 1 | *P. fluorescens* | 4.83±1.20<sup>a</sup> | 4.35±0.21<sup>b</sup> | 54.30±2.52<sup>b</sup> | 143.11±3.11<sup>c</sup> | 0.77±0.03<sup>a</sup> | 0.70±0.04<sup>a</sup> | 1.11±0.03<sup>a</sup> |
| PMH 1 | *B. subtilis* | 1.60±0.02<sup>b</sup> | 3.80±0.12<sup>bc</sup> | 70.51±0.51<sup>a</sup> | 125.33±3.52<sup>d</sup> | 0.77±0.04<sup>a</sup> | 0.80±0.01<sup>a</sup> | 1.53±0.01<sup>a</sup> |
| PMH 1 | *A. lipoferum* | 1.97±0.00<sup>b</sup> | 5.75±0.00<sup>a</sup> | 58.44±1.58<sup>c</sup> | 125.77±3.47<sup>e</sup> | 0.78±0.03<sup>a</sup> | 0.68±0.01<sup>a</sup> | 1.02±0.06<sup>a</sup> |
| PMH 1 | Control | 1.34±0.18<sup>c</sup> | 3.04±0.31<sup>d</sup> | 51.85±0.00<sup>d</sup> | 193.33±2.03<sup>a</sup> | 0.46±0.01<sup>b</sup> | 0.57±0.00<sup>b</sup> | 0.70±0.00<sup>b</sup> |
| PMH 10 | *P. putida* | 2.01±0.06<sup>b</sup> | 3.62±0.32<sup>c</sup> | 69.37±0.56<sup>b</sup> | 169.33±4.28<sup>b</sup> | 0.73±0.03<sup>a</sup> | 0.65±0.08<sup>a</sup> | 0.95±0.00<sup>a</sup> |
| PMH 10 | *P. fluorescens* | 2.12±0.06<sup>a</sup> | 4.79±0.12<sup>b</sup> | 68.22±1.02<sup>b</sup> | 112.00±5.55<sup>c</sup> | 0.77±0.05<sup>a</sup> | 0.67±0.12<sup>a</sup> | 1.03±0.02<sup>a</sup> |
| PMH 10 | *B. subtilis* | 2.35±0.12<sup>b</sup> | 3.73±0.12<sup>bc</sup> | 68.80±1.02<sup>a</sup> | 174.66±5.04<sup>d</sup> | 0.67±0.07<sup>a</sup> | 0.74±0.12<sup>a</sup> | 1.13±0.08<sup>a</sup> |
| PMH 10 | *A. lipoferum* | 1.69±0.18<sup>b</sup> | 3.04±0.12<sup>a</sup> | 64.11±0.87<sup>c</sup> | 120.00±0.00<sup>e</sup> | 0.69±0.06<sup>a</sup> | 0.71±0.13<sup>a</sup> | 0.97±0.04<sup>a</sup> |
| PMH 10 | Control | 0.95±0.26<sup>c</sup> | 2.43±0.19<sup>d</sup> | 55.73±0.00<sup>d</sup> | 230.22±20.44<sup>a</sup> | 0.46±0.00<sup>b</sup> | 0.40±0.15<sup>b</sup> | 0.47±0.00<sup>b</sup> |
| AXE* | *P. putida* | 1.98±0.02<sup>b</sup> | 5.56±0.11<sup>b</sup> | 65.12±1.04<sup>b</sup> | 214.66±2.66<sup>b</sup> | 0.67±0.00<sup>a</sup> | 0.76±0.10<sup>a</sup> | 0.98±0.02<sup>a</sup> |
| AXE* | *P. fluorescens* | 1.69±0.07<sup>a</sup> | 5.36±0.11<sup>b</sup> | 66.54±1.53<sup>b</sup> | 181.77±0.44<sup>c</sup> | 0.67±0.00<sup>a</sup> | 0.86±0.10<sup>a</sup> | 1.02±0.01<sup>a</sup> |
| AXE* | *B. subtilis* | 1.37±0.00<sup>b</sup> | 6.31±0.12<sup>bc</sup> | 65.42±0.98<sup>a</sup> | 97.33±0.00<sup>d</sup> | 0.69±0.00<sup>a</sup> | 0.76±0.10<sup>a</sup> | 1.06±0.02<sup>a</sup> |
| AXE* | *A. lipoferum* | 1.28±0.00<sup>b</sup> | 6.78±0.11<sup>a</sup> | 58.95±1.53<sup>c</sup> | 109.77±0.88<sup>e</sup> | 0.70±0.00<sup>a</sup> | 0.71±0.15<sup>a</sup> | 1.02±0.01<sup>a</sup> |
| AXE* | Control | 0.96±0.07<sup>c</sup> | 4.72±0.24<sup>d</sup> | 55.14±0.00<sup>d</sup> | 231.55±1.77<sup>a</sup> | 0.50±0.00<sup>b</sup> | 0.42±0.10<sup>b</sup> | 0.57±0.00<sup>b</sup> |

**Table 2.**
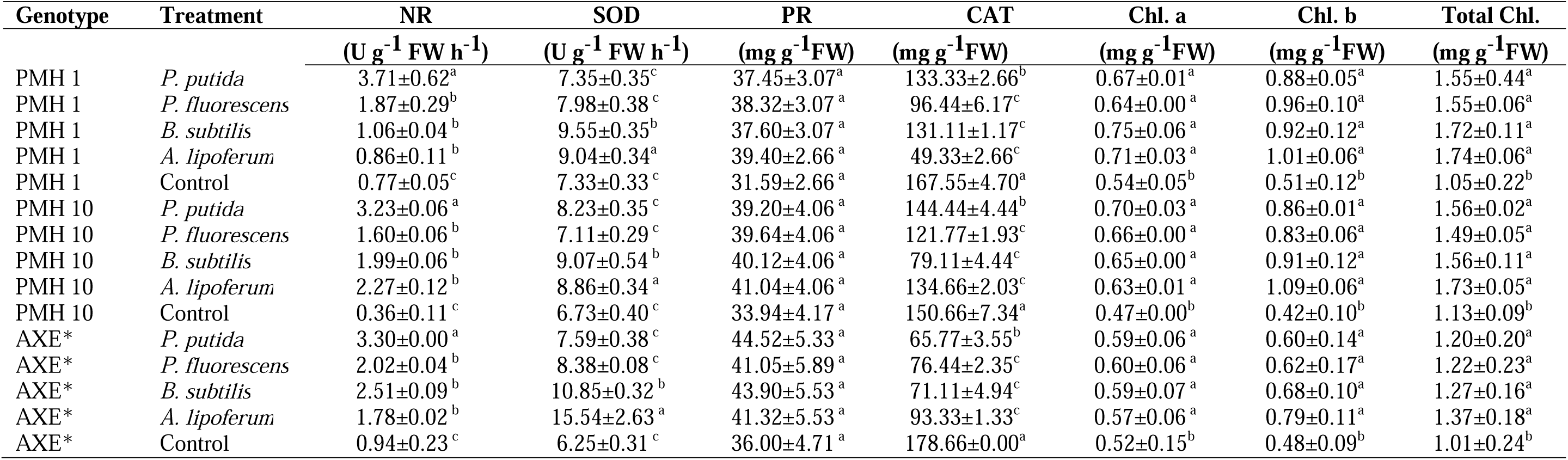
Nitrate reductase (NR)activity (U g^-1^ FW h^-1^), Superoxide dismutase (SOD) activity (U g^-1^ FW h^-1^), Protein content (PR) (mg g^-1^FW), Catalase activity (CAT) (mg g^-1^ FW), Chlorophyll a content (Chl. a) (mg g^-1^ FW), Chlorophyll b content (Chl. b) (mg g^-1^ FW), Total Chlorophyll content (Total Chl.) (mg g^-1^ FW) in the leaves of maize genotype treated with different bacterial strains and untreated control in vegetative stage in 2022. The values after plus-minus indicated deviation from the mean value. Means in the columns that have a common letter after them are not significantly different.

NR activity in the treated and untreated maize plants was also measured at the flowering stage. During 2021 (Table 3), the highest NR activity in the genotype AXE* was observed in the plants treated with *P. putida*, which recorded a value of 3.53 Ug^-1^FWh^-1^ against 1.99 Ug^-1^FWh^-1^ for control plants. In PMH1 highest NR activity was observed for the treatment *B. subtilis* with the value 3.76 Ug^-1^FWh^-1^ against 2.18 Ug^-1^FWh^-1^ for control and in PMH10 maximum NR activity was observed for plant treated with *A. lipoferum* that showed a value of 3.07 Ug^-1^FWh^-1^ against 2.32 Ug^-1^FWh^-1^ for control plants. Furthermore, during 2022 (Table 4) increased NR activity was found in plants inoculated with *P. fluorescens, A. lipoferum* and *P. putida* during blooming stage for genotypes AXE*, PMH 1, and PMH10, with increases of 132%(2.97 Ug^-1^FWh^-1^) (For control-1.28 Ug^-1^FWh^-1^), 121%(4.23 Ug^-1^FWh^-1^) (For control-1.91 Ug^-1^FWh^-1^), and 22%(3.66 Ug^-1^FWh^-1^) (For control-3.00 Ug^-1^FWh^-1^), respectively.

**Table 3.** Nitrate reductase (NR)activity (U g^-1^ FW h^-1^), Superoxide dismutase (SOD) activity (U g^-1^ FW h^-1^), Protein content (PR) (mg g^-1^FW), Catalase activity (CAT) (mg g^-1^ FW), Chlorophyll a content (Chl. a) (mg g^-1^ FW), Chlorophyll b content (Chl. b) (mg g^-1^ FW), Total Chlorophyll content (Total Chl.) (mg g^-1^ FW) in the leaves of maize genotype treated with different bacterial strains and untreated control in flowering stage in 2021. The values after plus-minus indicated deviation from the mean value. Means in the columns that have a common letter after them are not significantly different.

| Genotype | Treatment | NR<br>($\text{U g}^{-1} \text{FW h}^{-1}$) | SOD<br>($\text{U g}^{-1} \text{FW h}^{-1}$) | PR<br>($\text{mg g}^{-1} \text{FW}$) | CAT<br>($\text{mg g}^{-1} \text{FW}$) | Chl. a<br>($\text{mg g}^{-1} \text{FW}$) | Chl. b<br>($\text{mg g}^{-1} \text{FW}$) | Total Chl.<br>($\text{mg g}^{-1} \text{FW}$) |
| --- | --- | --- | --- | --- | --- | --- | --- | --- |
| PMH 1 | *P. putida* | 3.29±0.20<sup>a</sup> | 5.29±0.36<sup>ab</sup> | 84.01±0.51<sup>a</sup> | 92.88±4.44<sup>a</sup> | 0.74±0.01<sup>a</sup> | 0.40±0.01<sup>b</sup> | 1.15±0.02<sup>ab</sup> |
| PMH 1 | *P. fluorescens* | 2.57±0.37<sup>bc</sup> | 4.93±0.11<sup>b</sup> | 84.61±0.51<sup>a</sup> | 127.55±4.44<sup>a</sup> | 0.71±0.00<sup>a</sup> | 0.41±0.04<sup>a</sup> | 1.12±0.04<sup>a</sup> |
| PMH 1 | *B. subtilis* | 3.76±0.36<sup>ab</sup> | 4.88±0.14<sup>ab</sup> | 86.82±1.53<sup>a</sup> | 147.11±0.44<sup>b</sup> | 0.72±0.00<sup>a</sup> | 0.49±0.01<sup>c</sup> | 1.21±0.02<sup>c</sup> |
| PMH 1 | *A. lipoferum* | 2.56±0.09<sup>c</sup> | 6.16±0.15<sup>a</sup> | 83.54±0.00<sup>a</sup> | 141.33±0.00<sup>a</sup> | 0.73±0.01<sup>a</sup> | 0.39±0.01<sup>bc</sup> | 1.12±0.02<sup>bc</sup> |
| PMH 1 | Control | 2.18±0.20<sup>d</sup> | 3.93±0.03<sup>c</sup> | 57.82±4.09<sup>b</sup> | 180.44±0.88<sup>a</sup> | 0.51±0.00<sup>b</sup> | 0.14±0.00<sup>d</sup> | 0.67±0.01<sup>d</sup> |
| PMH 10 | *P. putida* | 2.89±0.20<sup>a</sup> | 4.19±0.18<sup>ab</sup> | 97.91±3.07<sup>a</sup> | 101.77±3.55<sup>a</sup> | 0.72±0.03<sup>a</sup> | 0.42±0.01<sup>b</sup> | 1.14±0.05<sup>ab</sup> |
| PMH 10 | *P. fluorescens* | 2.85±0.08<sup>bc</sup> | 4.13±0.09<sup>b</sup> | 101.90±5.22<sup>a</sup> | 92.44±0.88<sup>a</sup> | 0.70±0.00<sup>a</sup> | 0.74±0.03<sup>a</sup> | 1.44±0.03<sup>a</sup> |
| PMH 10 | *B. subtilis* | 2.93±0.13<sup>ab</sup> | 4.69±0.09<sup>ab</sup> | 98.48±3.38<sup>a</sup> | 153.33±0.00<sup>b</sup> | 0.73±0.01<sup>a</sup> | 0.30±0.02<sup>c</sup> | 1.04±0.04<sup>c</sup> |
| PMH 10 | *A. lipoferum* | 3.07±0.11<sup>c</sup> | 4.69±0.06<sup>a</sup> | 99.92±4.24<sup>a</sup> | 131.55±4.44<sup>a</sup> | 0.75±0.02<sup>a</sup> | 0.44±0.02<sup>bc</sup> | 1.20±0.05<sup>bc</sup> |
| PMH 10 | Control | 2.32±0.00<sup>d</sup> | 3.77±0.02<sup>c</sup> | 55.15±1.12<sup>b</sup> | 174.22±1.77<sup>a</sup> | 0.57±0.00<sup>b</sup> | 0.29±0.01<sup>d</sup> | 0.87±0.01<sup>d</sup> |
| AXE* | *P. putida* | 3.53±0.55<sup>a</sup> | 4.86±0.33<sup>ab</sup> | 102.76±5.83<sup>a</sup> | 141.33±0.00<sup>a</sup> | 0.69±0.00<sup>a</sup> | 0.60±0.03<sup>b</sup> | 1.30±0.02<sup>ab</sup> |
| AXE* | *P. fluorescens* | 2.59±0.37<sup>bc</sup> | 5.01±0.50<sup>b</sup> | 101.08±5.63<sup>a</sup> | 120.00±0.00<sup>a</sup> | 0.69±0.00<sup>a</sup> | 0.44±0.04<sup>a</sup> | 1.13±0.03<sup>a</sup> |
| AXE* | *Bacillus subtilis* | 2.52±0.13<sup>ab</sup> | 4.95±0.20<sup>ab</sup> | 104.04±6.09<sup>a</sup> | 77.33±0.00<sup>b</sup> | 0.67±0.00<sup>a</sup> | 0.42±0.06<sup>c</sup> | 1.09±0.05<sup>c</sup> |
| AXE* | *A. lipoferum* | 2.21±0.00<sup>c</sup> | 5.91±0.58<sup>a</sup> | 103.01±5.63<sup>a</sup> | 74.66±0.00<sup>a</sup> | 0.65±0.00<sup>a</sup> | 0.49±0.05<sup>bc</sup> | 1.14±0.04<sup>bc</sup> |
| AXE* | Control | 1.99±0.04<sup>d</sup> | 4.15±0.21<sup>c</sup> | 67.03±3.58<sup>b</sup> | 194.66±0.00<sup>a</sup> | 0.53±0.00<sup>b</sup> | 0.22±0.05<sup>d</sup> | 0.76±0.00<sup>d</sup> |

**Table 4.** Nitrate reductase (NR) activity (U g^-1^ FW h^-1^), Superoxide dismutase (SOD) activity (U g^-1^ FW h^-1^), Protein content (PR) (mg g^-1^FW), Catalase activity (CAT) (mg g^-1^ FW), Chlorophyll a content (Chl. a) (mg g^-1^ FW), Chlorophyll b content (Chl. b) (mg g^-1^ FW), Total Chlorophyll content (Total Chl.) (mg g^-1^ FW) in the leaves of maize genotype treated with different bacterial strains and untreated control in flowering stage in 2022. The values after plus-minus indicated deviation from the mean value. Means in the columns that have a common letter after them are not significantly different.

| Genotype | Treatment | NR | SOD | PR | CAT | Chl. a | Chl. b | Total Chl. |
| --- | --- | --- | --- | --- | --- | --- | --- | --- |
| | | ($\text{U g}^{-1} \text{FW h}^{-1}$) | ($\text{U g}^{-1} \text{FW h}^{-1}$) | ($\text{mg g}^{-1} \text{FW}$) | ($\text{mg g}^{-1} \text{FW}$) | ($\text{mg g}^{-1} \text{FW}$) | ($\text{mg g}^{-1} \text{FW}$) | ($\text{mg g}^{-1} \text{FW}$) |
| PMH 1 | *P. putida* | 1.99±0.67<sup>a</sup> | 12.37±3.72<sup>a</sup> | 41.40±3.44<sup>a</sup> | 171.11±0.44<sup>c</sup> | 0.68±0.06<sup>a</sup> | 0.68±0.10<sup>a</sup> | 1.13±0.16<sup>a</sup> |
| PMH 1 | *P. fluorescens* | 4.23±0.67<sup>a</sup> | 11.72±3.66<sup>a</sup> | 41.19±5.53<sup>a</sup> | 231.15±0.44<sup>d</sup> | 0.60±0.13<sup>ab</sup> | 0.43±0.11<sup>ab</sup> | 1.03±0.22<sup>ab</sup> |
| PMH 1 | *B. subtilis* | 2.56±0.69<sup>a</sup> | 12.17±3.85<sup>a</sup> | 43.31±3.17<sup>a</sup> | 272.88±1.77<sup>b</sup> | 0.69±0.07<sup>a</sup> | 0.56±0.10<sup>ab</sup> | 1.25±0.16<sup>a</sup> |
| PMH 1 | *A. lipoferum* | 1.95±0.68<sup>a</sup> | 11.43±3.88<sup>a</sup> | 43.51±3.35<sup>a</sup> | 270.66±4.00<sup>b</sup> | 0.69±0.07<sup>a</sup> | 0.54±0.11<sup>a</sup> | 1.23±0.16<sup>a</sup> |
| PMH 1 | Control | 1.91±0.71<sup>a</sup> | 8.09±1.13<sup>a</sup> | 38.00±3.01<sup>a</sup> | 322.22±3.55<sup>a</sup> | 0.56±0.01<sup>b</sup> | 0.34±0.15<sup>b</sup> | 0.90±0.13<sup>b</sup> |
| PMH 10 | *P. putida* | 3.66±0.65<sup>a</sup> | 7.62±3.68<sup>a</sup> | 43.24±4.60<sup>a</sup> | 172.44±1.77<sup>c</sup> | 0.65±0.07<sup>a</sup> | 0.63±0.12<sup>a</sup> | 1.29±0.19<sup>a</sup> |
| PMH 10 | *P. fluorescens* | 3.30±0.69<sup>a</sup> | 7.77±3.68<sup>a</sup> | 41.90±3.07<sup>a</sup> | 88.88±3.47<sup>d</sup> | 0.69±0.07<sup>ab</sup> | 0.49±0.12<sup>ab</sup> | 1.18±0.19<sup>ab</sup> |
| PMH 10 | *Bacillus subtilis* | 3.47±0.64<sup>a</sup> | 7.77±3.68<sup>a</sup> | 42.58±3.07<sup>a</sup> | 231.55±3.20<sup>b</sup> | 0.70±0.06<sup>a</sup> | 0.53±0.11<sup>ab</sup> | 1.14±0.18<sup>a</sup> |
| PMH 10 | *A. lipoferum* | 3.53±0.69<sup>a</sup> | 7.70±3.68<sup>a</sup> | 43.11±3.07<sup>a</sup> | 233.77±1.75<sup>b</sup> | 0.71±0.06<sup>a</sup> | 0.66±0.15<sup>a</sup> | 1.08±0.18<sup>a</sup> |
| PMH 10 | Control | 3.00±0.69<sup>a</sup> | 6.44±2.65<sup>a</sup> | 39.17±1.53<sup>a</sup> | 263.55±1.17<sup>a</sup> | 0.48±0.09<sup>b</sup> | 0.17±0.01<sup>b</sup> | 0.66±0.08<sup>b</sup> |
| AXE* | *P. putida* | 1.84±0.29<sup>a</sup> | 7.11±1.10<sup>a</sup> | 39.41±1.53<sup>a</sup> | 137.77±0.44<sup>c</sup> | 0.69±0.07<sup>a</sup> | 0.40±0.12<sup>a</sup> | 1.10±0.19<sup>a</sup> |
| AXE* | *P. fluorescens* | 1.71±0.13<sup>a</sup> | 7.48±1.10<sup>a</sup> | 38.75±1.53<sup>a</sup> | 132.88±3.11<sup>d</sup> | 0.57±0.06<sup>ab</sup> | 0.44±0.12<sup>ab</sup> | 1.02±0.19<sup>ab</sup> |
| AXE* | *B. subtilis* | 1.85±0.03<sup>a</sup> | 7.51±0.36<sup>a</sup> | 39.24±1.53<sup>a</sup> | 169.77±4.70<sup>b</sup> | 0.73±0.07<sup>a</sup> | 0.39±0.12<sup>ab</sup> | 1.06±0.18<sup>a</sup> |
| AXE* | *A. lipoferum* | 2.97±0.03<sup>a</sup> | 8.96±0.36<sup>a</sup> | 39.18±1.53<sup>a</sup> | 165.77±1.60<sup>b</sup> | 0.63±0.06<sup>a</sup> | 0.72±0.12<sup>a</sup> | 1.35±0.19<sup>a</sup> |
| AXE* | Control | 1.28±0.04<sup>a</sup> | 6.03±0.39<sup>a</sup> | 33.96±1.53<sup>a</sup> | 203.55±11.75<sup>a</sup> | 0.47±0.10<sup>b</sup> | 0.36±0.12<sup>b</sup> | 0.83±0.21<sup>b</sup> |

#### 3.2.2 Superoxide dismutase (SOD)

Enhanced SOD activity was found in all the genotypes under study during 2021 (Table 1). For AXE* and PMH1, the highest SOD activity was observed in plants treated with *A. lipoferum* with the recorded values of 66.54 Ug^-1^FWh^-1^ (55.14 Ug^-1^FWh^-1^ for control) and 70.51 Ug^-1^FWh^-1^ (51.75 Ug^-1^FWh^-1^ for control) respectively. However, in PMH10 maximum SOD activity was recorded in the plants treated with *P. fluorescens* with the value of 69.37 Ug^-1^FWh^-1^ in comparison to 55.73 Ug^-1^FWh^-1^ observed for control plants. However, during 2022 (Table 2) inoculation of AXE* with *A. lipoferum* shower the SOD activity of 15.54 Ug^-1^FWh^-1^ against 6.25 Ug^-1^FWh^-1^ for control plants. While in the genotypes PMH1 and PMH10 highest SOD activity was observed in the plants treated with *B. subtilis* with the observed values of 9.55 Ug^-1^FWh^-1^ against 7.33 Ug^-1^FWh^-1^ for control and 9.07 Ug^-1^FWh^-1^ against 6.73 Ug^-1^FWh^-1^ for control plants respectively.

In the year 2021, inoculation with *A. lipoferum* in AXE*, PMH1, and *A. lipoferum* and *B. subtilis* in PMH10 had the highest increases in SOD activity during the blooming period (Table 3). The rises for AXE* were 42% (5.91 Ug^-1^FWh^-1^ against the 4.15 Ug^-1^FWh^-1^for control), for PMH1 56% (6.16 Ug^-1^FWh^-1^ against 3.93 Ug^-1^FWh^-1^ of control) and 24% (4.69 Ug^-1^FWh^-1^ against 3.77 Ug^-1^FWh^-1^ for control) for PMH 10. Whereas in the year 2022 (Table 4), inoculation with *P. putida* and *A. lipoferum*, for genotypes PMH1 and AXE*, and *P. fluorescens* and *B. subtilis* for genotype PMH10, showed the highest increases in SOD activity where the observed values were 8.96 Ug^-1^FWh^-1^ against the control (6.03 Ug^-1^FWh^-^1) of AXE*, 12.37 Ug^-1^FWh^-1^ against the control (8.09 Ug^-1^FWh^-1^) for PMH1, and 7.77 Ug^-1^FWh^-1^ against the control (6.44 Ug^-1^FWh^-1^) for PMH 10.

#### 3.2.3 Protein Content

In the year 2021, at the vegetative stage (Table 1), the protein content was increased by 20.6% (66.54 mg g^-1^FW) when inoculated with *P. fluorescens* (For control- 55.14 mg g^-1^FW) for AXE*, by 35% (70.51 mg g^-1^FW) in the plants inoculated with *B. subtilis* (For control- 51.75 mg g^-1^FW) for PMH1, and by 24% (69.37 mg g^-1^FW) in plants inoculated with *P. putida* (For control- 55.73 mg g^-1^FW) for PMH 10. In year 2022, the protein content (Table 2) was increased by 23% (44.52 mg g^-1^FW) for the treatment *P. putida* (For control- 36.00 mg g^-1^FW) for AXE*, while for PMH1 and PMH10 maximum protein content was observed for the treatment *A. lipoferum* showing an increase of 24% (39.40 mg g^-1^FW) (For control- 31.59 mg g^-1^FW) for PMH1, and 20% (41.04 mg g^-1^FW) (For control- 33.94 mg g^-1^FW) for PMH 10.

During the flowering stage in the year 2021 (Table 3)*, B. subtilis* treated plants had the highest reported protein content for PMH1 and AXE* genotypes, while *P. fluorescens-*treated plants had the highest reported protein content for PMH10, with protein content increased by 55% (104.04 mg g^-1^FW) (For control- 67.63 mg g^-1^FW) for AXE*, 50% (86.82 mg g^-1^FW) (For control- 57.82 mg g^-1^FW) for PMH1, and 84% (101.90 mg g^-1^FW) (For control- 55.15 mg g^-1^FW) for PMH10. During the flowering stage in year 2022 (Table 4), *P. putida* recorded maximum protein content for genotype AXE* and PMH10, whereas *A. lipoferum* resulted in highest protein content for PMH1, with increase of 16% (39.41 mg g^-1^FW) (For control- 33.96 mg g^-1^FW) for AXE*, 14% (43.51 mg g^-1^FW) (For control- 38.00 mg g^-1^FW) for PMH1, and 10% (43.24 mg g^-1^FW) (For control- 39.17 mg g^-1^FW) for PMH 10.

#### 3.2.4 Chlorophyll a content

During the vegetative stage, in 2021 (Table 1), Chlorophyll a content increased by 40% (0.70 mg g^-1^FW against 0.50 mg g^-1^FW of control) for AXE*, 69% (0.78 mg g^-1^FW against 0.46 mg g^-1^FW for control) for PMH1, and 67% (0.77 mg g^-1^FW against 0.46 mg g^-1^FW for control) for PMH10. Inoculation with *A. lipoferum* showed the maximum chlorophyll content for AXE* and PMH1 while *P. fluorescens* recorded maximum chlorophyll content for PMH10. In the year 2022 (Table 2), AXE* had a 15% (0.60 mg g^-1^FW in comparison to 0.52 mg g^-1^FW in control plants) increase in chlorophyll content, PMH1 had a 38% (0.75 mg g^-1^FW in comparison to 0.54 mg g^-1^FW in control plants) increase, and PMH10 had a 58% (0.70 mg g^-1^FW in comparison to 0.47 mg g^-1^FW in control plants) increase. The treatments with *P. fluorescens* for AXE*, *B. subtilis* for PMH1, and *P. putida* for PMH10 observed an increase in chlorophyll content.

At the flowering stage, in the year 2021 (Table 3), The genotypes AXE*, PMH1, and PMH10 showed the maximum amount of chlorophyll content when inoculated with *P. putida* and *P. fluorescens* in AXE*, *P. putida* in PMH1 and *A. lipoferum* in PMH10, with an increment of 30%, 45% and 31% respectively for the three genotypes under study. However, during 2022 (Table 4), highest chlorophyll a content was observed for the treatments *B. subtilis* for AXE*, *A. lipoferum* and *B. subtilis* for PMH1, and *A. Lipoferum* for PMH10, with a rise of 45% (0.73 mg g^-1^FW against a value of 0.47 mg g^-1^FW in control plants) for AXE*, 23% (0.69 mg g^-1^FW against a value of 0.56 mg g^-1^FW of control) for PMH1, and 47% (0.71 mg g^-1^FW against a value of 0.48 mg g^-1^FW for control plants) of PMH10.

#### 3.2.5 Chlorophyll b content

The results obtained for chlorophyll b content at the vegetative stage during 2021 (Table 1), showed that chlorophyll b content increased from 0.42 mg g^-1^ FW found in control plants to 0.86 mg g^-1^ FW in the plants treated with *P. fluorescens* in AXE*, while in PMH1, it increased from 0.57 mg g^-1^ FW in control to 0.80 mg g^-1^ FW in plants treated with *B. subtilis* and in PMH10, it increased from 0.4 mg g^-1^ FW in control plants to 0.74 mg g^-1^ FW in plants treated with *B. subtilis*. During the year 2022 (Table 2), *A. lipoferum* was the most effective in increasing the chlorophyll b content across all genotypes. In the genotype AXE*, chlorophyll b content increased from 0.48 mg g^-1^ FW in control plants to 0.79 mg g^-1^ FW in treated plants. while in PMH1, it increased from 0.51 to 1.01 mg g^-1^ FW and in PMH10 from 0.42 to 1.09 mg g^-1^ FW in treated plants.

Similarly, at flowering stage in the year 2021 (Table 3), chlorophyll b content increased in the genotype AXE* from 0.22 mg g^-1^ FW in control to 0.60 mg g^-1^ FW in plants treated with *P. putida*, while in PMH1 the observed increase was from 0.14 to 0.49 mg g^-1^ FW in plants treated with *B. subtilis* and in PMH10, the observed increase was from 0.29 to 0.72 mg g^-1^ FW respectively. However, during the year 2022 (Table 4) *A. lipoferum* was effective for the genotypes AXE* and PMH10 showing an increase in chlorophyll b content from 0.36 to 0.72 mg g^-1^ FW in AXE* and from 0.48 to 0.71 mg g^-1^ FW in PMH10. While in PMH1, the most effective treatment was observed in *P. putida* showing an increase from 0.34 mg g^-1^ FW in control plants to 0.68 mg g^-1^ FW in treated plants.

#### 3.2.6 Total chlorophyll content

The results obtained for total chlorophyll content at the vegetative stage during 2021 (Table 1), showed that treatment of plants with *B. subtilis* was most effective in increasing total chlorophyll content in all three genotypes. In AXE*, total chlorophyll content increased from 0.57 to 1.06 mg g^-1^ FW, in PMH1 it increased from 0.70 to 1.53 mg g^-1^ FW and in PMH10, it increased from 0.47 to 1.13 mg g^-1^ FW. Whereas, during the year 2022 (Table 2), treatment with *A. lipoferum* was most effective in increasing the total chlorophyll content of treated plants of all three genotypes under study. In AXE*, total chlorophyll content increased from 1.01 to 1.37 mg g^-1^ FW, in PMH1, the increase in total chlorophyll content was from 1.05 mg g^-1^ FW in control plants to 1.74 mg g^-1^ FW in treated plants and PMH10, it increased from 1.13 to 1.73 mg g^-1^ FW.

At the flowering stage in the year 2021 (Table 3), there was a similar increase in the total chlorophyll content in the treated plants. For the genotype AXE* highest total chlorophyll content was observed in the plants treated with *P. putida,* with the observed values of 1.3 mg g^-1^ FW against 0.76 mg g^-1^ FW in control plants. While *B. subtilis* was most effective in increasing the chlorophyll in PMH1 (1.21 mg g^-1^ FW against 0.67 mg g^-1^ FW for control) and *P. fluorescens* was most effective in PMH10 (1.44 mg g^-1^ FW against 0.76 mg g^-1^ FW for control). Whereas in the year 2022 (Table 4), highest amount of total chlorophyll were recorded in the plants treated with *A. lipoferum* in AXE*, *B. subtilis* in PMH1, and *P. putida* in PMH10 with an increase of 62% (1.35 mg g^-1^ FW compared to 0.83 mg g^-1^ FW in control plants) for AXE*, 38% (1.25 mg g^-1^ FW compared to 0.90 mg g^-1^ FW in control plants) for PMH1 and 95% (1.29 mg g^-1^ FW compared to 0.66 mg g^-1^ FW in control plants) for PMH10.

#### 3.2.7 Catalase activity

Catalase activity was found to be reduced over both the years and for both stages. In the year 2021, at the vegetative stage (Table 1), for the genotype AXE*, the lowest catalase activity was observed in plants treated with *B. subtilis* (97.33 U mg^-1^FW) as compared to control (231.55 U mg^-1^FW). In PMH1, the lowest catalase activity was observed in plants treated with *P. fluorescens* (125.33 U mg^-1^FW) in comparison to control plants (193.33 U mg^-1^FW) and in PMH10 also, the lowest catalase activity was observed for the treatment *P. fluorescens* (112.0 U mg^-1^FW) in comparison to control plants (230.22 U mg^-1^FW). Whereas, during the year 2022 (Table 2), catalase activity was reduced by 171% (65.77 U mg^-1^FW) (for control-178.66 U mg^-1^FW) for AXE*, 73% (49.33 U mg^-1^FW) (for control-167.55 U mg^-1^FW) for PMH1, and 90% (79.11 U mg^-1^FW) (for control-150.66 U mg^-1^FW) for PMH10 inoculation with *P. putida, A. lipoferum,* and *B. subtilis* respectively.

However, at the flowering stage during 2021 (Table 3), inoculation with *A*. *lipoferum, P. putida,* and *P. fluorescens* experienced the maximum decline in catalase activity in the three genotypes respectively. For AXE*, catalase activity reduced by 160% (74.66 U mg^-1^FW against 194.66 U mg^-1^FW in control), while in PMH1, it declined by 44% (92.88U mg^-1^FW against 180.44 U mg^-1^FW in control plants), and in PMH10, it decreased by 88% (92.44 U mg^-1^FW against 174.22 U mg^-1^FW for control). Whereas, during 2022 (Table 4), inoculation of *P*. *fluorescens* for AXE* and PMH10, and *P. putida* for PMH1, showed the maximum decline in catalase activity with 53% (137.88 U mg^-1^FW compared to 203.55 U mg^-1^FW of control) for AXE*, 88% (171.11 U mg^-1^FW compared to 322.22 U mg^-1^FW of control) for PMH1, and 196% (88.88 U mg^-1^FW compared to 263.55 U mg^-1^FW of control) for PMH10.

### 3.3 Grain nutritional parameters

In order to ascertain the nutritional status of the grains their protein, carbohydrate, phytic acid and methionine content was also determined during both the years and data is presented in Tables 5 and 6 respectively.

**Table 5.** Protein content grain (mg g^-1^DW), carbohydrate content grain (mg g^-1^ DW), phytic acid grain (mg g^-1^DW) and methionine content (mg g^-1^ DW) of grain genotype treated with different bacterial strains and untreated control in harvesting stage in 2021. The values after plus-minus indicated deviation from the mean value. Means in the columns that have a common letter after them are not significantly different.

| Genotype | Treatment | Protein grain<br>(mg g<sup>-1</sup> DW) | Carbohydrate grain<br>(mg g<sup>-1</sup> DW) | Phytic acid<br>(mg g<sup>-1</sup> DW) | Methionine grain<br>(mg g<sup>-1</sup> DW) |
| --- | --- | --- | --- | --- | --- |
| PMH 1 | *P. putida* | 71.18±1.53<sup>a</sup> | 2.64±0.03<sup>bc</sup> | 4.06±0.20<sup>b</sup> | 16.54±2.95<sup>a</sup> |
| PMH 1 | *P. fluorescens* | 79.80±1.53<sup>a</sup> | 2.80±0.01<sup>c</sup> | 4.19±0.21<sup>b</sup> | 16.74±2.95<sup>a</sup> |
| PMH 1 | *B. subtilis* | 80.33±1.53<sup>a</sup> | 3.11±0.00<sup>a</sup> | 4.03±0.21<sup>b</sup> | 17.20±2.95<sup>a</sup> |
| PMH 1 | *A. lipoferum* | 82.61±2.58<sup>a</sup> | 2.86±0.01<sup>b</sup> | 6.26±1.15<sup>b</sup> | 16.91±2.95<sup>a</sup> |
| PMH 1 | Control | 62.00±0.42<sup>b</sup> | 1.92±0.02<sup>d</sup> | 9.69±1.27<sup>a</sup> | 10.91±1.97<sup>b</sup> |
| PMH 10 | *P. putida* | 79.22±0.20<sup>a</sup> | 2.98±0.00<sup>bc</sup> | 4.27±0.01<sup>b</sup> | 20.15±2.62<sup>a</sup> |
| PMH 10 | *P. fluorescens* | 76.42±0.88<sup>a</sup> | 3.03±0.00<sup>c</sup> | 4.66±0.00<sup>b</sup> | 19.57±1.97<sup>a</sup> |
| PMH 10 | *B. subtilis* | 76.72±1.30<sup>a</sup> | 3.03±0.00<sup>a</sup> | 4.37±0.08<sup>b</sup> | 20.32±1.97<sup>a</sup> |
| PMH 10 | *A. lipoferum* | 80.41±0.46<sup>a</sup> | 3.01±0.00<sup>b</sup> | 4.37±0.08<sup>b</sup> | 19.58±1.97<sup>a</sup> |
| PMH 10 | Control | 71.84±0.69<sup>b</sup> | 2.51±0.05<sup>d</sup> | 7.86±0.12<sup>a</sup> | 14.22±0.98<sup>b</sup> |
| AXE* | *P. putida* | 50.86±2.56<sup>a</sup> | 3.22±0.02<sup>bc</sup> | 4.66±0.00<sup>b</sup> | 8.35±0.98<sup>a</sup> |
| AXE* | *P. fluorescens* | 45.91±3.07<sup>a</sup> | 2.80±0.08<sup>c</sup> | 4.38±0.42<sup>b</sup> | 8.50±0.98<sup>a</sup> |
| AXE* | *B. subtilis* | 42.06±4.71<sup>a</sup> | 3.12±0.01<sup>a</sup> | 4.09±0.42<sup>b</sup> | 10.76±0.19<sup>a</sup> |
| AXE* | *A. lipoferum* | 45.67±4.15<sup>a</sup> | 3.02±0.09<sup>b</sup> | 4.51±0.42<sup>b</sup> | 9.16±0.16<sup>a</sup> |
| AXE* | Control | 36.21±3.09<sup>b</sup> | 1.58±0.11<sup>d</sup> | 7.28±0.29<sup>a</sup> | 4.33±0.19<sup>b</sup> |

**Table 6.** Protein content grain (mg g^-1^DW), carbohydrate content (mg g^-1^ DW) phytic acid (mg g^-1^DW) and methionine content grain (mg g ^-1^ DW) genotype treated with different bacterial strains and untreated control in harvesting stage in 2022. The values after plus-minus indicated deviation from the mean value. Means in the columns that have a common letter after them are not significantly different.

| Genotype | Treatment | Protein grain<br>($\text{mg g}^{-1}\text{DW}$) | Carbohydrate grain<br>($\text{mg g}^{-1}\text{DW}$) | Phytic acid<br>($\text{mg g}^{-1}\text{DW}$) | Methionine grain<br>($\text{mg g}^{-1}\text{DW}$) |
| --- | --- | --- | --- | --- | --- |
| PMH 1 | *P. putida* | 6.28±1.47<sup>a</sup> | 3.14±0.07<sup>a</sup> | 2.34±0.01<sup>b</sup> | 9.56±0.94<sup>ab</sup> |
| PMH 1 | *P. fluorescens* | 8.07±1.36<sup>a</sup> | 2.57±0.02<sup>d</sup> | 2.30±0.02<sup>b</sup> | 9.38±0.92<sup>ab</sup> |
| PMH 1 | *B. subtilis* | 8.71±1.17<sup>a</sup> | 3.12±0.02<sup>b</sup> | 2.17±0.13<sup>b</sup> | 10.16±0.92<sup>a</sup> |
| PMH 1 | *A. lipoferum* | 9.09±1.33<sup>a</sup> | 2.94±0.00<sup>c</sup> | 2.21±0.02<sup>b</sup> | 10.37±0.91<sup>a</sup> |
| PMH 1 | Control | 5.37±1.05<sup>a</sup> | 1.78±0.03<sup>e</sup> | 2.67±0.12<sup>a</sup> | 7.97±0.31<sup>b</sup> |
| PMH 10 | *P. putida* | 9.86±0.12<sup>a</sup> | 3.09±0.00<sup>a</sup> | 2.36±0.09<sup>b</sup> | 11.34±0.85<sup>ab</sup> |
| PMH 10 | *P. fluorescens* | 8.35±1.40<sup>a</sup> | 3.01±0.00<sup>d</sup> | 2.20±0.23<sup>b</sup> | 11.72±0.87<sup>ab</sup> |
| PMH 10 | *B. subtilis* | 7.91±1.35<sup>a</sup> | 3.02±0.00<sup>b</sup> | 2.24±0.13<sup>b</sup> | 12.54±0.91<sup>a</sup> |
| PMH 10 | *A. lipoferum* | 7.70±1.35<sup>a</sup> | 3.06±0.00<sup>c</sup> | 2.25±0.08<sup>b</sup> | 12.78±0.90<sup>a</sup> |
| PMH 10 | Control | 6.82±1.46<sup>a</sup> | 2.21±0.00<sup>e</sup> | 2.69±0.01<sup>a</sup> | 10.48±0.57<sup>b</sup> |
| AXE* | *P. putida* | 7.39±1.42<sup>a</sup> | 3.19±0.00<sup>a</sup> | 2.43±0.06<sup>b</sup> | 6.85±0.95<sup>ab</sup> |
| AXE* | *P. fluorescens* | 7.60±1.40<sup>a</sup> | 2.79±0.01<sup>d</sup> | 2.38±0.05<sup>b</sup> | 7.00±0.94<sup>ab</sup> |
| AXE* | *B. subtilis* | 7.36±1.41<sup>a</sup> | 3.02±0.00<sup>b</sup> | 2.36±0.02<sup>b</sup> | 8.16±0.89<sup>a</sup> |
| AXE* | *A. lipoferum* | 7.75±1.40<sup>a</sup> | 2.94±0.00<sup>c</sup> | 2.33±0.03<sup>b</sup> | 7.98±0.87<sup>a</sup> |
| AXE* | Control | 6.89±1.48<sup>a</sup> | 1.75±0.03<sup>e</sup> | 2.73±0.12 | 6.19±0.90<sup>b</sup> |

#### 3.3.1 Grain protein content

In year 2021, protein content of cereal grains (Table 5) increased by 40% in AXE* plants treated with *P. putida*, 33% in PMH1 and 11% in PMH10 plants inoculated with *A. lipoferum* in comparison to the control plants. During the year 2022 (Table 6), protein content in the seed kernels of AXE* increased by 12% and in PMH1 it increased by 69% and the most responsive treatment was *A. lipoferum*, while in grains of PMH10, protein content was increased by 44% in the plants inoculated with *P. putida* in comparison to the control plants.

#### 3.3.2 Carbohydrate content

In the year 2021, carbohydrate content of grains (Table 5) increased by 97% for AXE*, 61% for PMH1 and 20% for PMH10 in comparison to the control plants. The most responsive treatment for improving carbohydrate content was *P. putida* in AXE* and *B. subtilis* in PMH1, while for the genotype PMH10, treatment with *P. fluorescens* and *B. subtilis* was equally effective. During the year 2022 (Table 6), carbohydrate content increased by 82% for AXE*, 76% in PMH1 and 39% in PMH10, in comparison to control plants. Treatment of plants with *P. putida* was beneficial in improving the carbohydrate content of the grains in all three genotypes.

#### 3.3.3 Phytic acid

In the year 2021, a decrease in phytic acid content was observed in all the grains of all three genotypes and all the bacterial treatments (Table 5). In the genotype AXE*, there was 77% decline in the phytic acid content in grains obtained from plants treated with *B. subtilis*, while in PMH1, the decline was 140% in the grains obtained from the treatment *B. subtilis* and there was a decline of 84% in phytic acid content in grains of PMH10 obtained from plants treated with *P. putida* in comparison to the control plants. However, during the year 2022 (Table 6), the reduction in phytic acid for AXE* was 17% in the grains of plants treated with *A. lipoferum*, for PMH1, it was 23% in the grains obtained from plants treated with *P. fluorescens* and in PMH10, it was 22% in the treatment *B. subtilis* in comparison to the control plants.

#### 3.3.4 Methionine content

In the year 2021 (Table 5), methionine content in the grains obtained from the plants treated with *B. subtilis* showed a maximum increase in all three genotypes under study. In AXE*, methionine content increased by 57.6%, in PMH1 by 42.8% and in PMH10 by 148% in comparison to the control plants. Also, during the year 2022, treatment with *B. subtilis* was most effective in increasing the methionine content of grains in AXE* showing an increase of 31.8%, whereas, *A. lipoferum* was most effective in increasing the methionine content in grains of PMH1 and PMH10 with the recorded increase of 30% and 20% respectively in comparison to the control plants.

### 3.4 Yield attributes

The ultimate aim of any experiment where grain crops are concerned is productivity. Hence, data related to crop productivity including the parameters like total cob weight, cob weight (without bract), 100 grain weight, yield/m^2^ and projected yield/ha has been determined and the data for the years 2021 and 2022 is presented in Figure 2-6. In the year 2021, data suggested that inoculation with PGPR on three genotypes had considerably larger weights of cob with bract (Figure 2) than uninoculated plants. The highest increases in weight of cob with bract were seen in genotypes PMH1 and AXE* inoculated with *A. lipoferum*, while PMH10 plants treated with *B. subtilis*, respectively with an increase observed in AXE* by 52.2%, PMH1 by 45%, and PMH10 by 53%. Whereas, during 2022 *B. subtilis* was most effective in increasing the weight of cob with bracts in case of genotypes AXE* and PMH1 showing an increment of 42% and 32% respectively. While for the genotype PMH10, the treatment with *A. lipoferum* was most effective in improving the cob weight with bract showing an increase of 35% in comparison to the control plants.

**Figure 2.**
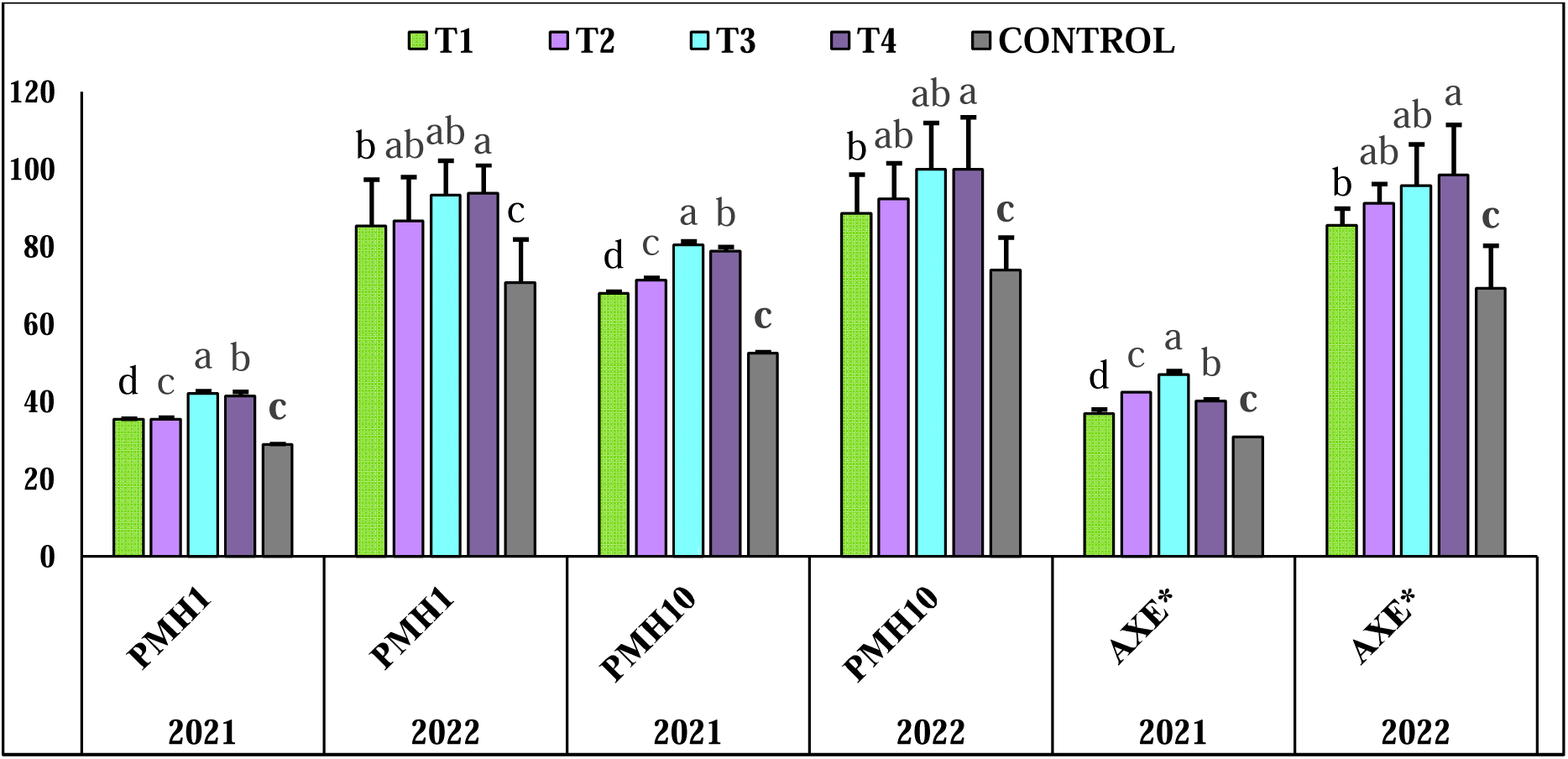
Cob weight (g) of the three maize genotypes AXE*, PMH1 and PMH10 treated with *P. putida, P. fluorescens, A. lipoferum*, *B. subtilis* and control during the planting seasons 2021 and 2022.

All three genotypes which were inoculated with PGPR had considerably increased weight of cob without bract as compared to non-inoculated plants(Figure 3). The weight of the cob without bract increased by 60% for AXE*, 71.44% for PMH1, and 55% for PMH10 and data revealed that inoculation with *A. lipoferum* for PMH1 and *B. subtilis* for PMH10 and AXE* genotypes showed the biggest gains in cob weight without bract. While during 2022, inoculated plants showed an increase of 54% for AXE*, 53% for PMH1, and 60% for PMH10 and the highest increases in weight of cob without bract were seen in *A. lipoferum* inoculated AXE*, PMH1, and *B. subtilis* treated plants of genotype PMH10.

**Figure 3.**
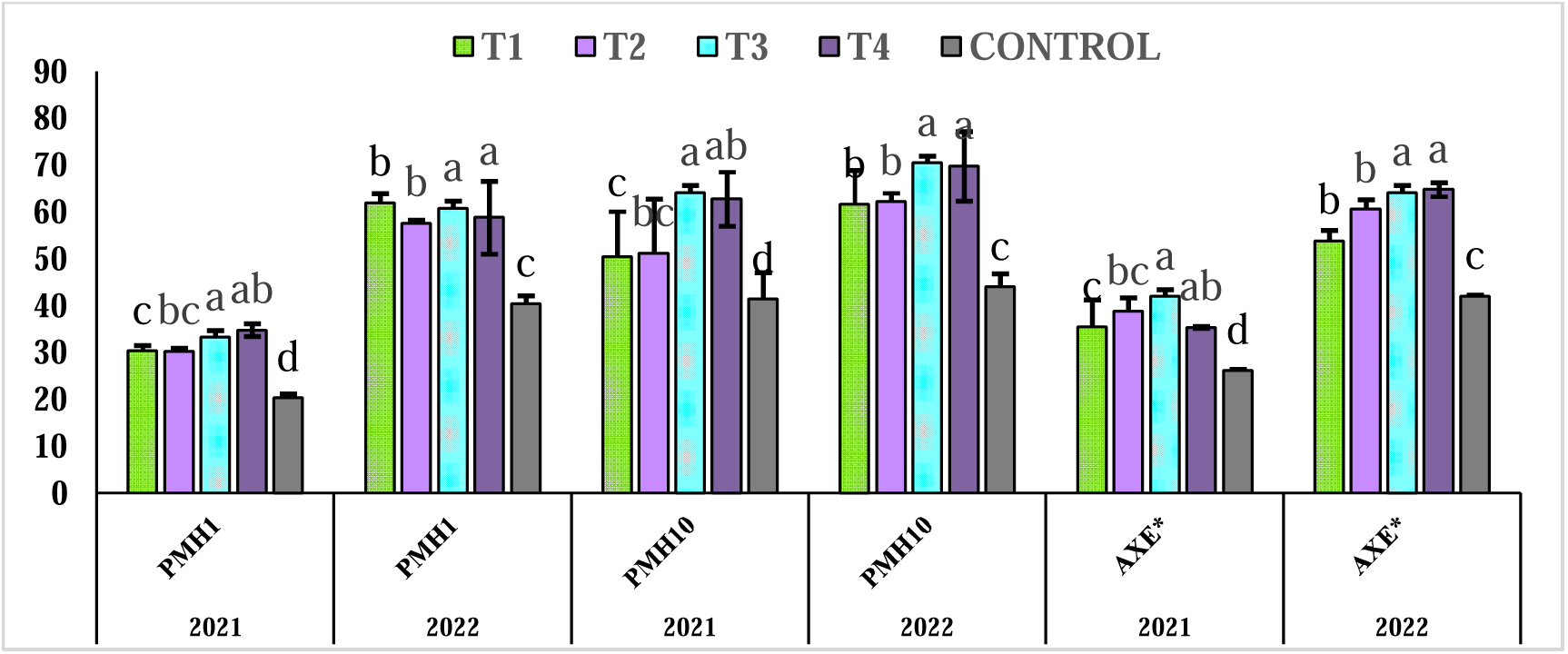
Cob weight without bract (g) of the three maize genotypes AXE*, PMH1 and PMH10 treated with *P. putida, P. fluorescens, A. lipoferum*, *B. subtilis* and control during the planting seasons 2021 and 2022.

For all three genotypes, the findings demonstrated that inoculation of plants with PGPR showed much weighted 100 grains compared to the control plants (Figure 4). There was an increase of 14% in grain weight of 100 for AXE*, 25% for PMH1, and 32% for PMH10, with the maximum 100 grain weight reported for inoculation with *B. subtilis* in AXE* and PMH1 genotypes, and *P. putida* in PMH10. Also in the year 2022, all three genotypes showed that PGPR-treated plants produced significantly more weighted 100 grains than control plants. There was a 14% increase in weight of 100 grains for AXE*, 25.9% increase for PMH1, and 22.8% increase for PMH10, and recorded highest grain weight per 100 in *A. lipoferum* for PMH1 and AXE*, and *B. subtilis* for PMH10.

**Figure 4.**
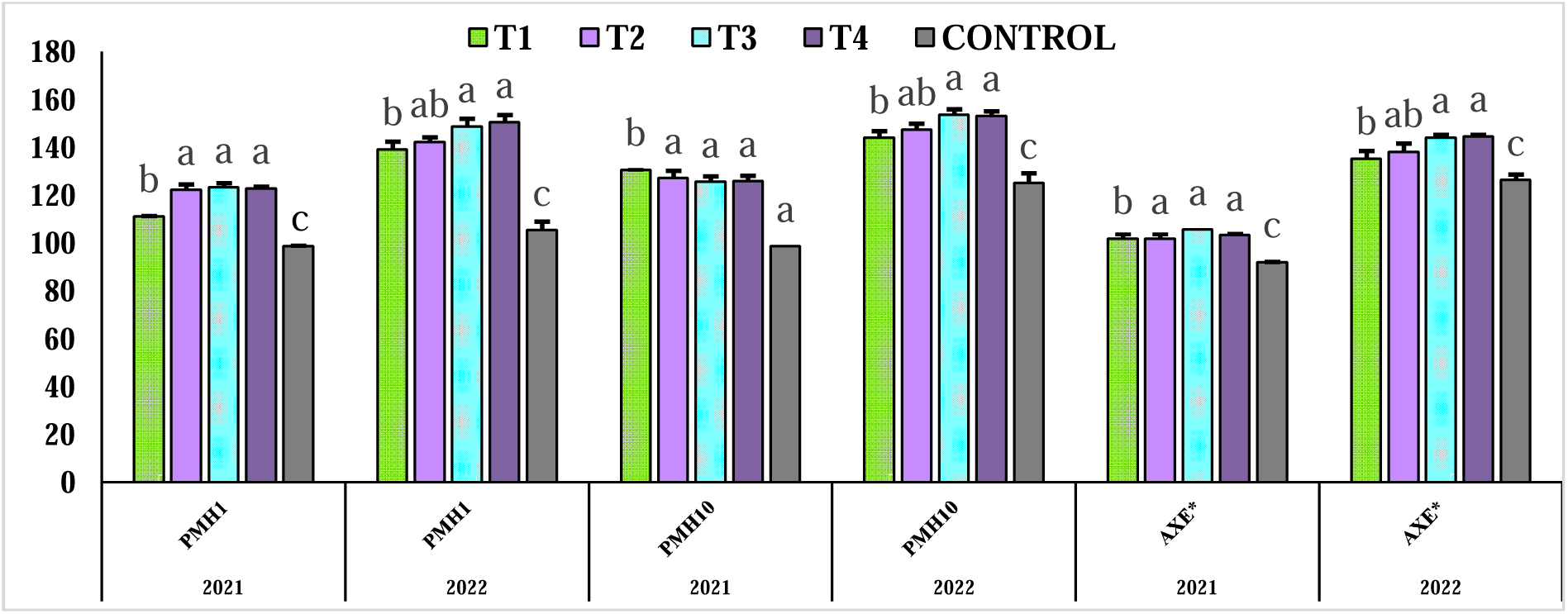
Hundred grains weight (g) of the three maize genotypes AXE*, PMH1 and PMH10 treated with *P. putida, P. fluorescens, A. lipoferum*, *B. subtilis* and control during the planting seasons 2021 and 2022.

The statistical analysis showed that three genotypes inoculated with PGPR showed better performance as compared to non-inoculated plants in terms of yield per m^2^ (Figure 5) with an increase of 58% for AXE*, 41% for PMH1, and 47% for PMH10 and plants treated with *B. subtilis* producing the highest yield per m^2^. During the year 2022, increase in yield per m^2^ was 17.6% for AXE*, 14% for PMH1, and 18% for PMH10, and inoculation with *B. subtilis* for PMH10 as well as inoculation with *A. lipoferum* in PMH1 and AXE* reported highest yield per m^2^.

**Figure 5.**
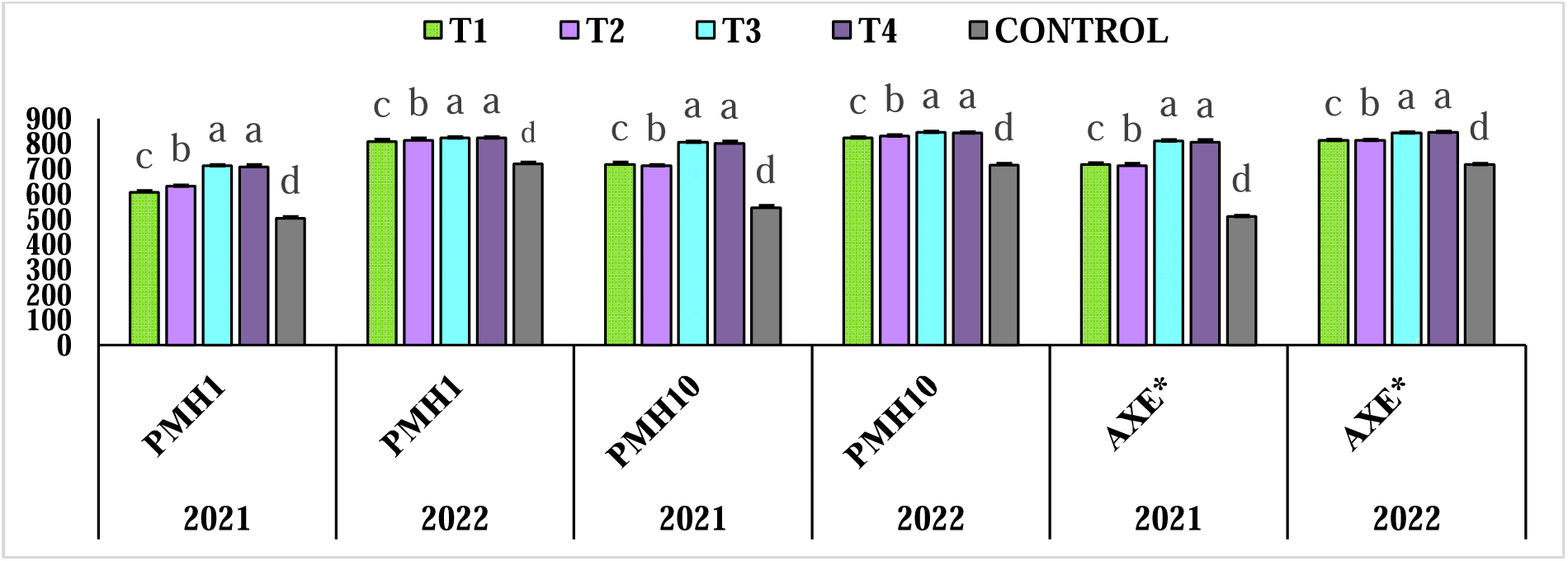
Yield (g m^-2^) of the three maize genotypes AXE*, PMH1 and PMH10 treated with *P. putida, P. fluorescens, A. lipoferum*, *B. subtilis* and control during the planting seasons 2021 and 2022.

Similarly, findings showed that inoculation with PGPR outperforms control plants in terms of yield per hector (Figure 6), with an increase of 58% for AXE*, 41% for PMH1, and 47% for PMH10 and plants treated with *B. subtilis* reporting the highest yield per hector. Similarly, during the year 2022, findings showed plants inoculated with PGPR outperform control plants in terms of yield per hector with an increase of17.6% for AXE*, 14% for PMH1, and 18% for PMH10 and inoculation with *B. subtilis* for PMH10 as well as inoculation with *A. lipoferum* for AXE* and PMH1 reporting the highest yield per hector.

**Figure 6.**
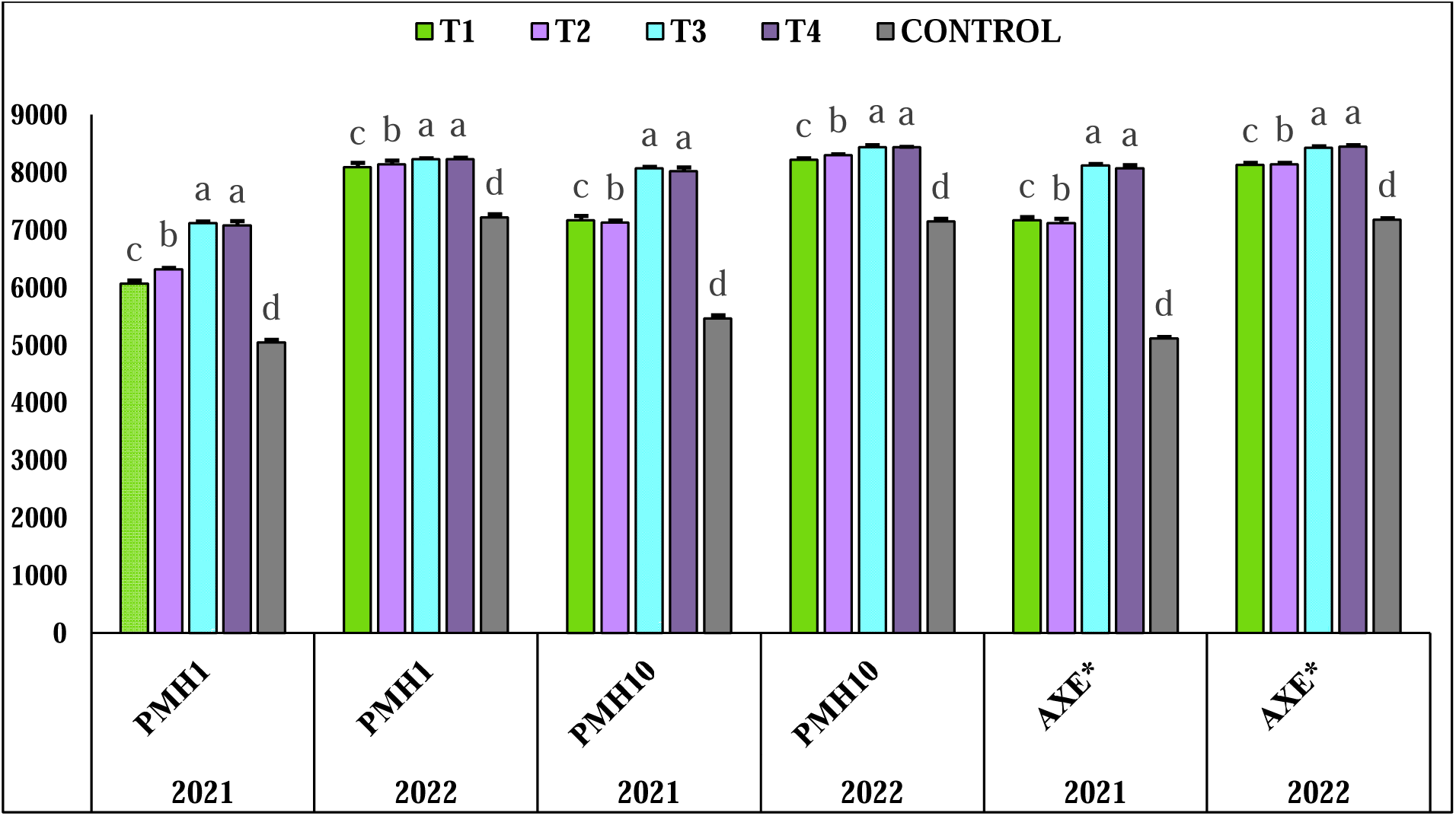
Yield (g hectare^-1^) of the three maize genotypes AXE*, PMH1 and PMH10 treated with *P. putida, P. fluorescens, A. lipoferum*, *B. subtilis* and control during the planting seasons 2021 and 2022.

## 4. Discussion

The present study found that different bacterial inoculations in three varieties of maize improved the numbers of leaves, fresh weight and dry weight of treated plant when compared to non-inoculated plants continuously over both the years. PGPR strains *B. subtilis* and *A. lipoferum* have shown a more pronounced effect as compared to other bacteria under study. Danish et al. (2019) reported comparable outcomes for maize, where they stated *Azospirillum* inoculation in plants leads to a remarkable escalation for a number of leaves, fresh weight, and dry weight of plants by 53%, 126.4% and 147.2%, respectively. Similar reports were given by Chaudhary et al. (2021), where they stated that plants inoculated with *Bacillus spp*. displayed enhanced fresh and dry weight.

PGPR strains *B. subtilis* and *P. putida* have shown a more pronounced effect on NR activity as compared to other bacteria under study. Both these bacteria have previously been shown to improve nitrate fascination, allowing plants to make maximum consumption of the nitrogen source (Cervantes-Vázquez et al., 2021; Huang et al., 2021), which advanced to improvement in seedling root development; increase in nitric oxide and auxins contents and hence the result obtained. According to Lee et al. (2020) for plants to adapt to difficult circumstances, nitrate-reducing enzyme activity is crucial, and *B. subtilis* L1 inoculation results in a significant increase in NR activity, as compared to the genotype that did not receive any bacterial inoculation. According to Kumar et al. (2019), PGPR inoculation boosted NR activity in pigeonpea.

According to the current investigation, application of bacteria to three genotypes increased SOD activity in comparison to non-inoculated seeds at different periods. PGPR strain *A. liopferum* has shown a more pronounced effect as compared to other bacteria under study. Previous research has demonstrated that these bacteria boost antioxidant defence system by reducing ROS, phytohormone modulation, and transcriptional control of host stress-responsive genes (Akhtar et al., 2021), which can lead to improvement in stress-tolerance ability in plants. Ali et al. (2020) got an identical outcome, claiming a noteworthy growth in SOD activity of 23-36% in maize treated with PGPR when compared to non-inoculated plants.

Liu et al. (2021b) suggested the first step in increasing maize’s dietary value is to enhance its level of protein throughout its leaves as well as grains. Data from this study demonstrated that bacterial inoculation raised protein content in three genotypes compared with controls in different periods. All PGPR strains have shown a positive effect on inoculated plants under study. All bacteria have previously been shown to fix atmospheric nitrogen to facilitate N absorption as well as the synthesis of critical components (Pagnani et al., 2020), increased concentration of which can lead to improvement in plant physiological and metabolic activities and hence the results obtained. (Abdel Latef et al., 2020) observed comparable outcomes in maize, indicating that plants inoculated with *A. lipoferum* had a greater protein content than control plants. In the same way, Ashraf et al. (2019), Chaudhary et al. (2021) and Yasmin et al. (2017) reported that inoculation with PGPR especially *Bacillus* sp. and *Pseudomonas* sp. boosted protein content by 30.43% in wheat, 24% in maize, and in drought stressed maize plants, inoculation with *Pseudomonas* sp. led to 85% increase in protein content while on inoculation with *B. pumilus* increase was 87%, significantly higher than control plants.

The data showed that, at various stages, inoculation with bacteria in three genotypes improved chlorophyll concentration in comparison to control plant. All PGPR strains have shown a positive effect on inoculated plants under study. Abdel Latef et al. (2020) observed inoculation with *A. lipoferum* increased chlorophyll a and b content by 26% and 19% respectively and Shabaan et al. (2022) reported an increase in chlorophyll a content by 35% and in chlorophyll b content by 33% in maize plants treated with PGPRs.

The data showed that, after bacterial inoculation, all three genotypes of maize had higher glucose contents than the control plants. PGPR strain *B. subtilis* has shown a more pronounced effect as compared to other bacteria under study. Pandey et al. (2018) found inoculation with *Bacillus* increases *Amaranthus hypochondriacus* intake of vital nutrients in the soil, lending credence to the idea that the increase in chemical components is driven by *Bacilli* strains. A comparable result was also reported by Misra and Chauhan (2020) and Chaudhary et al. (2021) which is, inoculation with *bacillus* sp. showed an increase in carb content with 35.98% in maize.

In order to lessen CAT activity and the negative effects of drought stress, rhizobacteria are used. PGPR strains *P. putida* and *P. fluorescens* have shown a more pronounced effect as compared to other bacteria under study. Misra and Chauhan (2020) reported in maize plants treatment with *Bacillus* spp. displayed reduced CAT activity by 40.70% and Azeem et al. (2022) also reported comparable results with inoculation with different strains of *Bacillus* spp. decreases antioxidant activity by 62.96%.

Phytic acid is found in plant seeds and acts as an antinutrient. PGPR strain *B. subtilis* has shown a more pronounced effect as compared to other bacteria under study. Ghorbani Nasrabadi et al. (2023) reported decreased phytic acid activity by 1.65% to 95.2% in maize seeds inoculated with PGPR and these findings werealso supported by Singh et al. (2018) who had similar results in plants inoculated with PGPR.

Methionine is one of the most vital amino-acid needed for the synthesis of sulphur and glutathione, a free radical scavenger. PGPR strain *B. subtilis* has shown a more pronounced effect as compared to other bacteria under study. This report is also supported by Pandey et al. (2018) who reported a 47.68% increase in methionine content in *Amaranths hypochondriacus* plants treated with *B. sibtilis.* Similarly, Kaur et al. (2020) observed a 38.8% rise in methionine content in rice plants treated by PGPR species like *Pseudomonas* and *Bacillus*.

The results showed that PGPR inoculation increases agricultural output while favouring plant growth and development. PGPR strains *B. subtilis* and *A. lipoferum* have shown a more pronounced effect as compared to other bacteria under study. Both these bacteria have been shown to produce ACC deaminase and ethylene (Ali et al., 2020; Czarnes et al., 2020) that can improve plant growth and hence the results obtained. Akhtar et al. (2018) observed that inoculation with PGPR resulted in a higher fresh cob weight with bract (107.45 g cob^-1^), with gains of 52.69%, 49.4%, and 47.98% and combined application of several strains boosted cob weight by up to 40.87% over the non-inoculated control. Similarly, Latta and Eskin (1980) observed that gain in the weight of cob with husk by 79.33% as well as an increase in the weight of cob without husk by 36.33% with the inoculation of PGPR combinations in maize. According to Nezarat and Gholami (2009) weight per 100 grains increased by 44% with the inoculation of *Azospirillum* and *Pseudomonas* spp. The results of Chaudhary et al. (2021) and Siddaiah et al. (2018) were comparable as all inoculated maize plants had improved crop qualities due to elevated biochemical characteristics of maize plants, which culminated in a higher yield. The current results demonstrated that bacterial inoculation enhanced the yield per m^2^ and yield per hectare of the maize crops when compared to the control. The results demonstrated PGPR inoculation promoted the plant’s average weight in dry matter and showed a beneficial effect on increase in yield. Kumar et al. (2019) reported a comparable outcome in crops of maize inoculated with PGPR and possessing the greatest yield per hector of 8.6 ton/ha inoculated with *Pseudomonas* spp. and *B. subtilis.* Erdemci (2020) found similar result with *P. fluorescens* inoculated lentil had higher grain yield as compared to un-inoculated seeds by 20.3%.

## 5. Conclusion

The present study suggested that the application of PGPR in maize seedlings enriched development, metabolism and crop yield. The amounts of essential enzymes and nutrient quality of maize plants were all increased by the usage of PGPR, along with development, growth and grain yield. As an outcome, applying rhizobacteria to stimulate plant development on maize makes sense; additional investigation needs to focus on this notion. *A. lipoferum* and *B. subtilis*, in particular, showed optimistic results on the development of the crop. We suggest that *A. lipoferum* and *B. subtilis* are PGPR species exclusive to the maize plant. We also suggest that the application of PGPRs holds immense potential for improving the productivity of maize as well as other crops and future research needs to be focussed on the identification of species-specific beneficial bacteria and their judicial use to improve agricultural sustainability.

## Acknowledgement

This research was supported by the Basic Science Research Program through the National Research Foundation of Korea funded by the Ministry of Education (2020R1I1A3054816).The authors are also thankful to the Agricultural Research Organization (ARO-Volcani Center, Israel) for research support. The authors are also thankful to the editors and anonymous reviewers for their critical comments to improve the manuscript. We apologize to those researchers whose relevant research publications are not cited in this manuscript due to the space limitation in the present form.

## Authors Contributions

SS, RR, VM, AM and AS conceptualized the work, designed methodology, analysed data, project administration, and draft – review and editing; SS, HK, RK, NJ and YHA analysed the data and wrote the draft manuscript; SS and HK collected the data and assisted in draft writing and analysis; VK and MG reviewed and edited the manuscript; RR, AM and YHA assisted for research funding; All authors reviewed and edited the manuscript and agreed to the published version of the manuscript.

## Data Availability Statement

Will be made available on demand.

---

**Figure S1.**
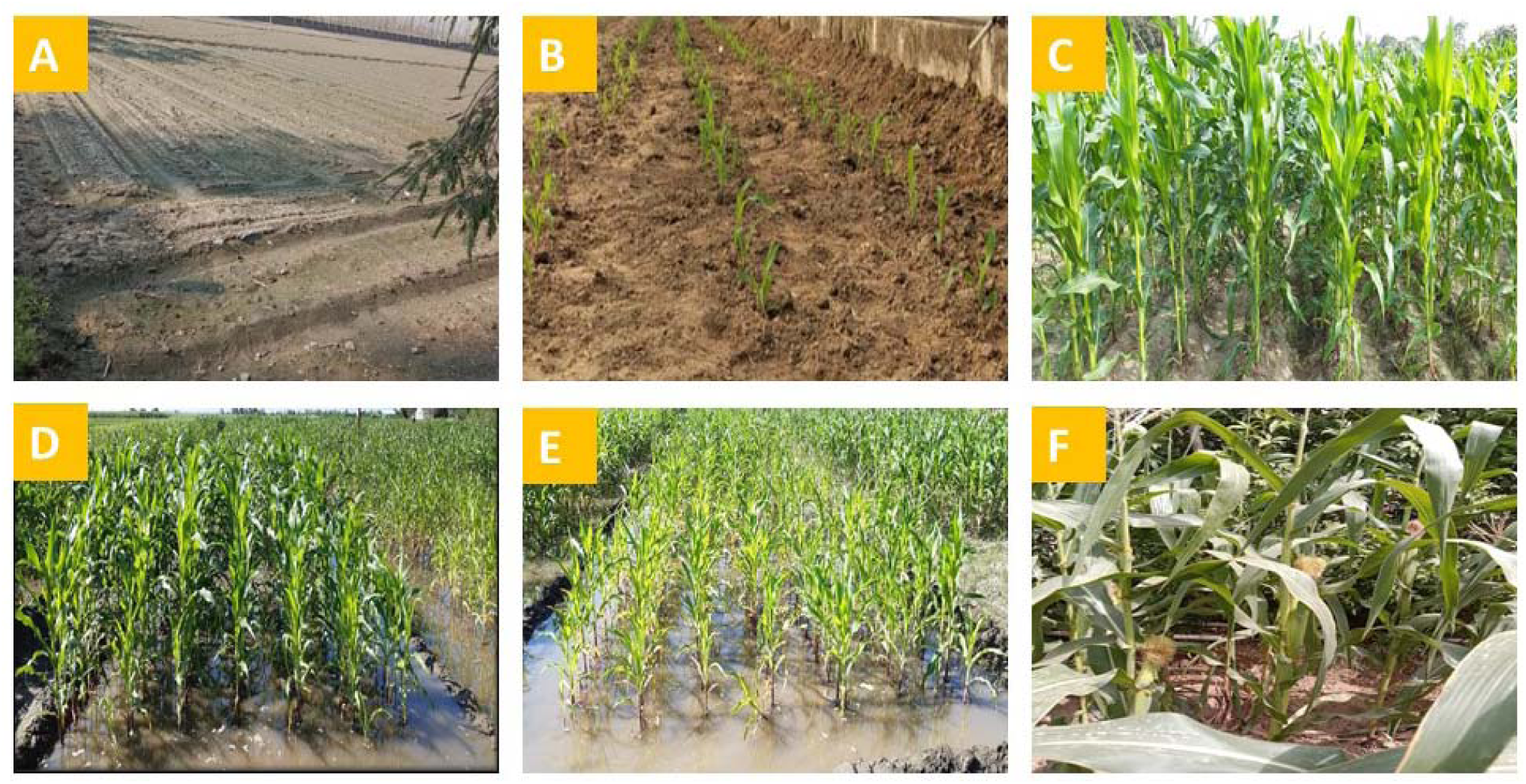
Cultivation of maize plants for the present investigation. (A) field preparation, (B) germinated maize seedlings, (C) Vegetative growth stage of maize plants, (D-E) irrigation of maize plots, (F) flowering in the maize crop.

